# A biaryl-linked tripeptide from *Planomonospora* leads to widespread class of minimal RiPP gene clusters

**DOI:** 10.1101/2020.07.21.214643

**Authors:** Mitja M. Zdouc, Mohammad M. Alanjary, Guadalupe S. Zarazúa, Sonia I. Maffioli, Max Crüsemann, Marnix H. Medema, Stefano Donadio, Margherita Sosio

**Affiliations:** Naicons Srl., Viale Ortles 22/4, 20139 Milano; Swammerdam Institute for Life Sciences, University of Amsterdam, Science Park 904, 1098 XH, Amsterdam, The Netherlands; Bioinformatics Group, Wageningen University, Droevendaalsesteeg 1, 6708 PB, Wageningen, The Netherlands; Institut für Pharmazeutische Biologie, Rheinische Friedrich-Wilhelms-Universität, Nuβallee 6, 53115 Bonn, Germany

## Abstract

Microbial natural products impress by their bioactivity, structural diversity and ingenious biosynthesis. While screening the rare actinobacterial genus *Planomonospora,* cyclopeptides **1A** and **1B** were discovered, featuring an unusual Tyr-His biaryl-bridging across a tripeptide scaffold, with the sequences *N*-acetyl-Tyr-Tyr-His (**1A**) and *N*-acetyl-Tyr-Phe-His (**1B**). Genome analysis of the **1A** producing strain pointed towards a ribosomal synthesis of **1A**, from a pentapeptide precursor encoded by the tiny 18-nucleotide gene *bycA,* to our knowledge the smallest gene ever reported. Further, biaryl instalment is performed by the closely linked gene *bycB,* encoding a cytochrome P450 monooxygenase. Biosynthesis of **1A** was confirmed by heterologous production in *Streptomyces,* yielding the mature product. Bioinformatic analysis of related cytochrome P450 monooxygenases indicated that they constitute a widespread family of pathways, associated to 5-aa coding sequences in approximately 200 (actino)bacterial genomes, all with potential for a biaryl linkage between amino acids 1 and 3. We propose the name biarylicins for this newly discovered family of RiPPs.

---

Microbial natural products impress not only for their potent bioactivity, but also for their chemical diversity and originality. Structurally novel compounds have a high chance of interacting with their cognate targets with mechanisms of action different from known compounds. ^1^ In recent years, ribosomally synthesized and post-translationally modified peptides (RiPPs) have drawn attention due to their intricate, three-dimensional structures and wide distribution in bacteria. RiPP biosynthetic gene clusters (BGC) are compact, encompassing only a precursor peptide, separated in a N-terminal leader and a C-terminal core peptide. After ribosomal translation, the precursor peptide is modified by dedicated enzymes, with consecutive cleavage of the leader peptide and release of the mature core peptide.^2^ Although several bioinformatic tools have been developed for the automatic recognition of RiPP BGCs in microbial genomes, they are usually based on recognition elements such as leader peptide or processing enzymes.

In the search for new microbial secondary metabolites, we investigated 72 strains belonging to the rare actinomycete genus Planomonospora by extensive, HR-ESI-LC-MS/MS based metabolome mining of extracts.^4^ Molecular networking analysis revealed a group of related features (see Figures S1-2), observed in samples from four phylogenetically related strains, with 16S rRNA gene sequences having 99.5 to 99.6% identity with *Planomonospora algeriensis* PM3.^4^ One strain, *Planomonospora* sp. ID82291, produced a compound – **1A** – with an *m/z* 522.198 [M+H]^+^ (calculated molecular formula C_26_H_28_N_5_O_7_), while the other three strains, with different 16S rRNA gene sequences, produced a compound – **1B** – with an *m/z* 506.203 [M+H]^+^ (calculated molecular formula C_26_H_28_N_5_O_6_). **1A** and **1B** showed UV absorption maxima at 270 and 315 nm, the latter indicating an unusual chromophore.

*Planomonospora* sp. ID82291 was selected for further investigation. Extraction and purification of **1A** proved difficult due to high solubility of **1A** in water. Eventually, 5 mg pure **1A** were isolated from 15L of cultivation broth. Extensive 1D and 2D NMR analysis indicated an acetylated short peptide. A first set of experiments, performed in DMSO-*dh*, allowed the observation of three aromatic spin systems containing CHα and CH2β consistent with two tyrosine and one histidine residues. The correlations observed in a NOESY-experiment suggested the sequence *N*-acetyl-Tyr-Tyr-His, but with an additional unsaturation, which could not be assigned due to overlapping aromatic signals. A second set of experiments in CD_3_OD allowed the complete assignment of the non-exchangeable protons of **1A**. TOCSY and HSQC experiments showed that the histidine residue lacked its characteristic aromatic proton at position 5”, but had a quaternary carbon at 142 ppm. In an HMBC experiment, a diagnostic cross-correlation was observed between both CH-5 and CH-8 and C5”, consistent with a C-C bond between C6 and C5”. The existence of the cross-link was confirmed after hydrolysis of **1A** (see Figure S3), which lead to the release of a compound with *m/z* 335 [M+H]^+^ at 4.4 minutes, corresponding to the expected carbon-carbon linked tyrosine and histidine, and a compound with *m/z* 480 [M+H]^+^ at 5.74 minutes, consistent with the loss of the acetyl-group. The *m/z* 335 [M+H]^+^ compound retained the UV maximum at 315nm, as expected for the biaryl chromophore.

Compound **1B**, produced by the other three *Planomonospora* isolates, was deduced to be *N*-acetyl-Tyr-Phe-His, based on the similarity of the tandem mass fragmentation of **1A** and **1B**, with a neutral loss corresponding to phenylalanine (−147.07 Da) being the only difference between the fragmentation spectra (see Figure S4). Since genomic analysis indicates a ribosomal synthesis (see below), **1A** is predicted to consist of L-amino acids only, similarly to K13 (**2**) and OF4949 (**3**).

We could only identify one literature precedent for natural products with a tyrosine-histidine biaryl bridge: aciculitins, cyclic nonapeptides decorated with glycolipids from the marine sponge *Aciculites orientalis,* contain a carbon-carbon bond between tyrosine and histidine aromatic moieties, but involving C6 of histidine instead of C5 as in **1A**. Interestingly, aciculitins, which lack additional aromatic residues, show UV-absorption maxima at 270 and 310 nm, matching those of **1A** and **1B**.^5^ A few metabolites have been described that contain a cross-linked *N*-acetylated tripeptide: K-13 (**2**) from *Micromonospora halophytica* ssp. *exilisia* with a Tyr-Tyr-Tyr sequence and an ether bond between the oxygen of Tyr-3 and C6 of Tyr-1;^6,7^ OF-4949 (**3**) from *Peni-cillinum rugulosum* OF4949, which is identical to **2** except for asparagine in place of the central tyrosine;^8^ and pseudosporamide (**4**) from *Pseudosporangium* sp. RD062863, reported while writing this manuscript, with a Tyr-Pro-Trp sequence and a carbon-carbon bond between C6 of tyrosine and C-9 of tryptophan.^9^ Of note, compounds **2**, **3** and **4** lack an absorption maximum at 315nm. It should also be noted that compound **2**, **3** and **4** have all been established to consist only of L-amino acids. Among the **1A**-related metabolites, **2** and **3** have been reported to inhibit the proteases angiotensin-converting enzyme (ACE) and aminopeptidase B, respectively, with apparent strict specificity, since **3** is not an ACE inhibitor. We were unable to observe inhibition of different proteases by **1A** (data not shown), suggesting that the amino acid sequence and/or ring size can strongly influence activity. **4** was reported to have weak cytotoxic activity.^9^

The production of **1A** increased with time (Figure S5) and, when the strain was grown in D_2_O-supplemented medium, **1A** was found to be extensively labelled with deuterium (Figure S6), indicating that its formation requires *de novo* amino acid synthesis.

We are unaware of any biosynthetic studies on **2**, **3** or **4**. To get insights into **1A** biosynthesis, we analysed the 7.58-Mbp genome of *Planomonospora* sp. ID82291^4^ for the presence of antiSMASH-predicted NRPS or RiPP biosynthetic gene clusters (BGCs),^10^ likely to specify a Tyr-Tyr-His peptide. However, this search proved unsuccessful, suggesting that **1A** was formed by a BGC not recognized by the antiSMASH search tools. Searching the 6-frame translation of the ID82291 genome led to a single plausible candidate: a short open reading frame (ORF) encoding the pentapeptide MRYYH, preceded by a Shine-Dalgarno ribosomal binding site (RBS) and followed by PLM4_2056, encoding a cytochrome P450 monooxygenase (see Figure 3A). The proposed BGC for **1A**, encoding the two genes designated *bycA* and *bycO* (see below), is similar to the recently reported BGC for tryptorubin (**5**), a hexapeptide containing carbon-carbon and carbon-nitrogen bonds between aromatic amino acid residues.^11,12^ Also for **5**, the BGC encodes just one cytochrome P450 enzyme and a precursor peptide, the latter however containing a 22-aa leader peptide. Similarly, the recently reported BGC for cittilin (**6**), a cross-linked YIYY tetrapeptide from *Myxococcus xanthus,* encodes a 27-aa precursor peptide, with an adjacent cytochrome P450 enzyme, a methyltransferase and a distant endopeptidase for cleavage. ^13^ Cytochrome P450 monooxygenases are known to install cross-links between aromatic residues of peptides, as e.g. in the NRPS-generated glycopeptides. ^14^ These precedents made the BGC depicted in Figure 3A as the likely candidate for **1A** biosynthesis. Apart from a possible regulator, no additional genes that could partake in **1A** biosynthesis are present in close vicinity.

**Figure 1:**
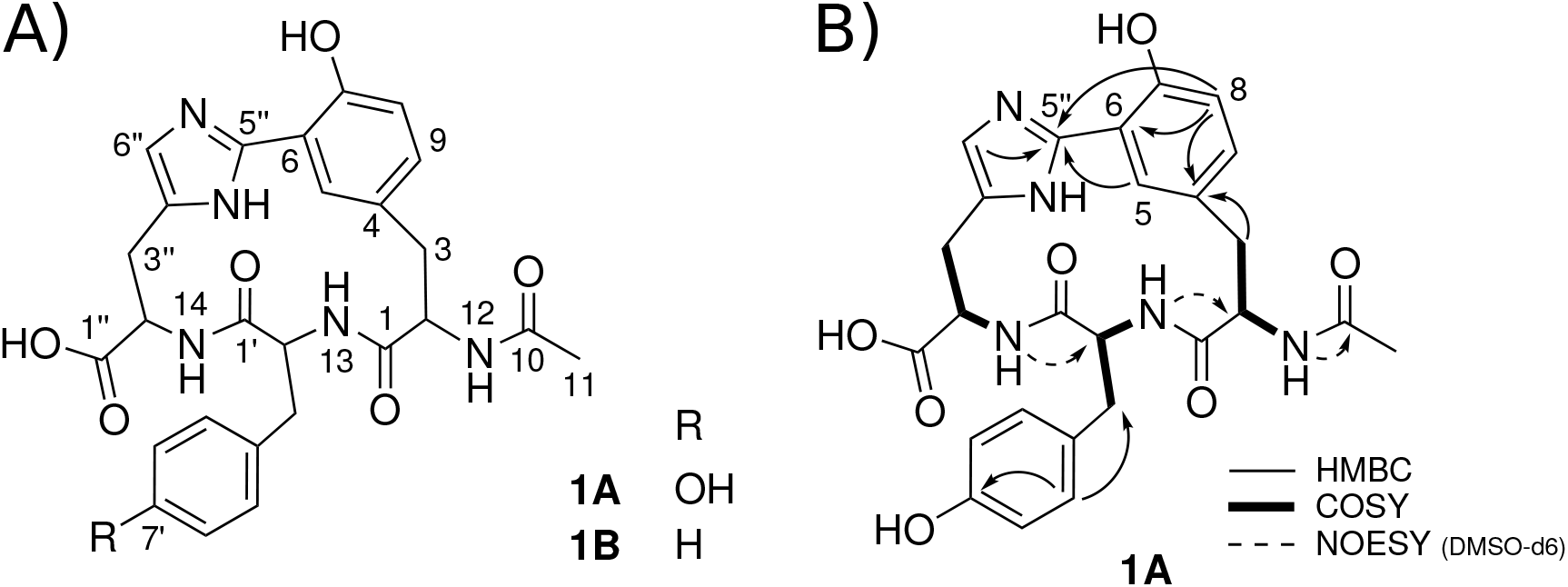
(A) structures of **1A** and **1B**. (B) key 2D NMR correlations for **1A**.

**Figure 2:**
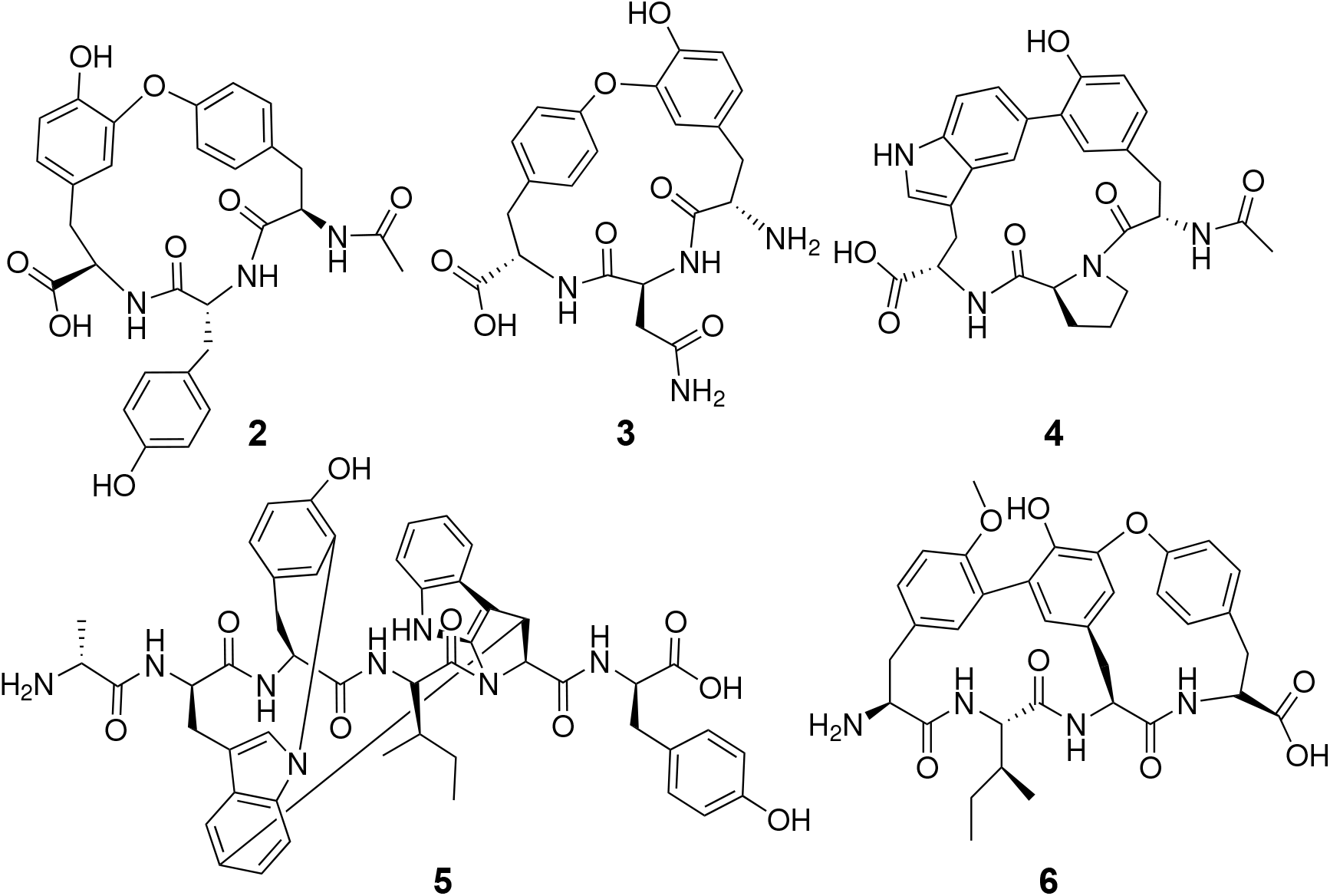
Related metabolites: K-13 (**2**), OF-4949 (**3**), pseudosporamide (**4**), tryptorubin (**5**) and cittilin A (**6**)

**Figure 3:**
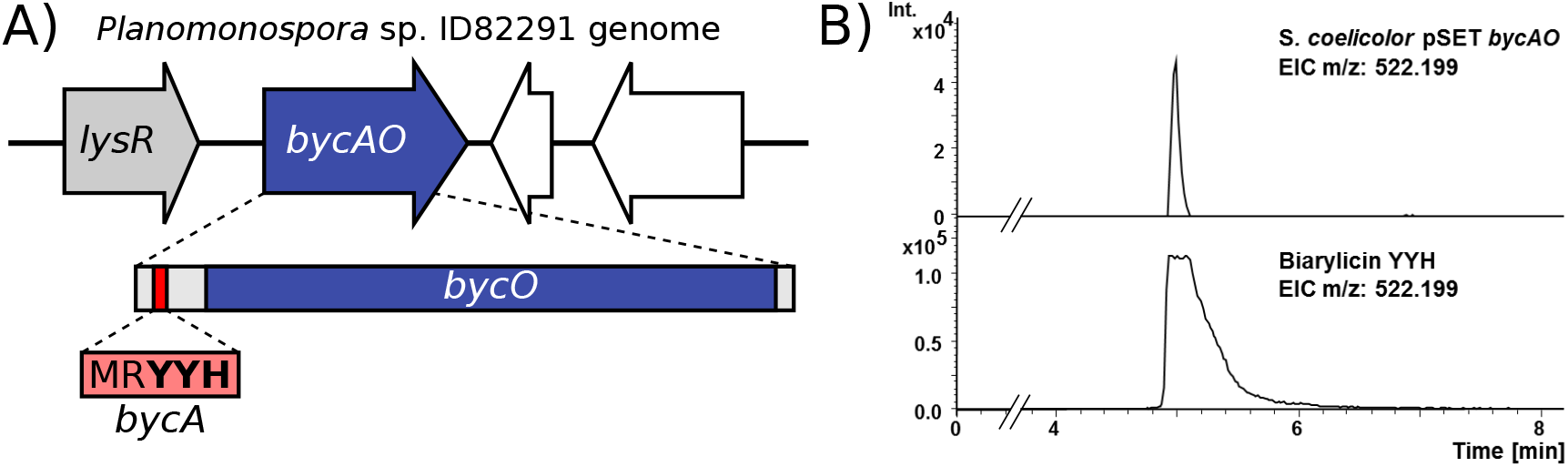
(A) visualization of the region for **1A** biosynthesis. *bycA* encodes the 5-aa precursor peptide, while *bycO* encodes the cytochrome P450 monooxygenase PLM4_2056. A *lysR* family transcriptional regulator is in vicinity of *bycAO.* (B) extracted ion chromatograms (EIC) for biarylicin YYH from heterologous production in *Streptomyces coelicolor* M1152 (top) and an authentic standard of biarylicin YYH, isolated from *Planomonospora* sp. ID82291 (bottom).

To prove that biarylicin production is indeed encoded by these two genes, we cloned a 1303 base pair sequence containing *bycA* and *bycO* into pSET152 and conjugated the plasmid into the expression host *Streptomyces coelicolor* M1152. Cultivation of *S. coelicolor* pSET *byc*AO, followed by HP20 extraction and HPLC-MS/MS analysis, resulted in the detection of a peak with a *m/z* 522.199, having the same UV spectrum, retention time and MS^2^ spectrum as **1A** (Figure 3B, Figure S7). This unambiguously confirms the heterologous production of mature **1A** by the minimal gene cluster *bycAO*, the shortest RiPP BGC so far reported. The biarylicin core BGC therefore consists only of two genes: the precursor peptide-encoding gene *bycA* and *bycO*, encoding the P450 monooxygenase. To the best of our knowledge, *bycA* is also the smallest gene ever reported across the tree of life, encompassing only 18 bp (including stop-codon), in comparison to mccA (21 bp), involved in microcin-biosynthesis, previously considered the smallest gene. ^15^ Since heterologous expression of *bycAO* in *Streptomyces coelicolor* resulted in the mature product, we infer that **1A** biosynthesis recruits one or more generic enzymes, involved in trimming the additional amino acid(s) at the *N*-terminus and in *N*-acetylation. Of note, the existence of cluster-encoded proteases in RiPP BGCs from *Actinobacteria* seems to be the exception.^16^

Further, we reasoned that the cytochrome P450 monooxygenase PLM4_2056 would be selective in installing a biaryl cross-link on short peptides, and that it could belong to a distinct phylogenetic branch of this enzyme family. If the cognate pentapeptide-encoding gene was closely linked to the cytochrome monooxygenase gene, then one could easily identify additional BGCs related to the one depicted in Figure 3A. We thus conducted a search for analogous small CDSs in the flanking regions of genes related to PLM4_2056. Assuming structural conservation of the carbon-carbon bond, we restricted our search to pentapeptides with tyrosine at position 3 and tyrosine/histidine at position 5. From approximately 3,300 PLM_2056-related sequences, CDS encoding the specified peptides were identified in approximately 200 genomes, the majority from *Actinobacteria*. To increase our confidence in the hits, we filtered the detected CDS for a Shine-Dalgarno motive 6-8 bp upstream of the peptide sequence, as indicated in the phylogenetic trees in Figures 4 and 5. Interestingly, such a ribosomal binding site was only detected in sequences from actinobacteria, where it occurred in vicinity of approximately half of the detected peptides (indicated by filled circles in Figures 4 and 5). Furthermore, this clade was seen to have the only examples of replicate motifs (denoted “2x” in Figure 5). To get insight into the amino acid distribution, detected peptides were aligned using the program MUSCLE. Peptides with a preceding ribosomal binding site showed less variation, with a strong preference for a basic amino acid at position 2, in comparison to the global alignment (see Figures 4 and S8). Intriguingly, the diversity of genomic loci in which PLM_2056 homologues were found together with byc-like motifs suggests that these BGCs encode the production of a much wider diversity of molecules with distinct chemical modifications, as suggested by the co-localization of methyltransferases, sulfotransferases, ATP-grasp enzymes and other tailoring enzymes in these loci (Figure 5). Together, this represents a treasure trove for further discovery efforts in this class.

**Figure 4:**
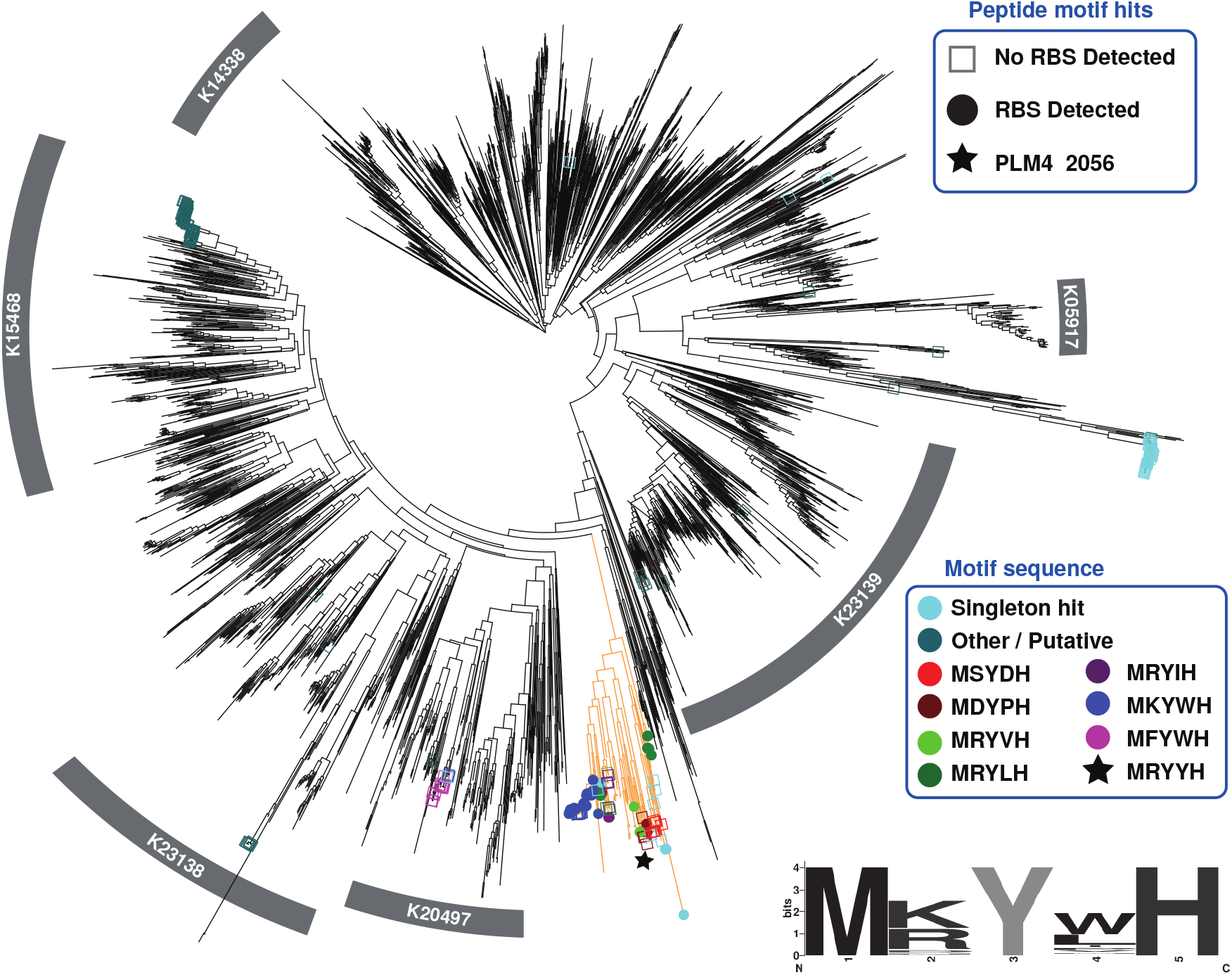
Overview phylogeny of cytochrome p450s from a variety of bacterial genera, used in this study. KEGG orthology IDs in grey boxes highlight selected functional groupings (K20497: methyl-branched lipid omega-hydroxylase, K23138: Family CYP109, K23139: Family CYP110, K05917: Steroid biosynthesis, K15468: Polyketide biosynthesis, K14338: Fatty acid degradation). Positive matches for pentapeptide motif hits are indicated as circles, where filled ones visualize a fully validated Shine-Dalgarno (SD) ribosomal binding site (RBS), while for empty ones, a SD sequence was not detected. Putative singleton hits are marked in light or dark cyan if the same motive is not seen in other p450s, with <95% amino acid identity, or if the hit overlaps with the P450 coding region, respectively. Identical motives correspond to the same color shown in the legend. P450s likely associated with biarylicins are highlighted in orange. This clade also contains the *bycAO* sequence marked with a filled star. Consensus sequences of all motifs with validated RBS sites are illustrated with the sequence logo at the bottom right. More annotations and interactive tree can be accessed at: https://itol.embl.de/tree/21312710611256691590408100

**Figure 5:**
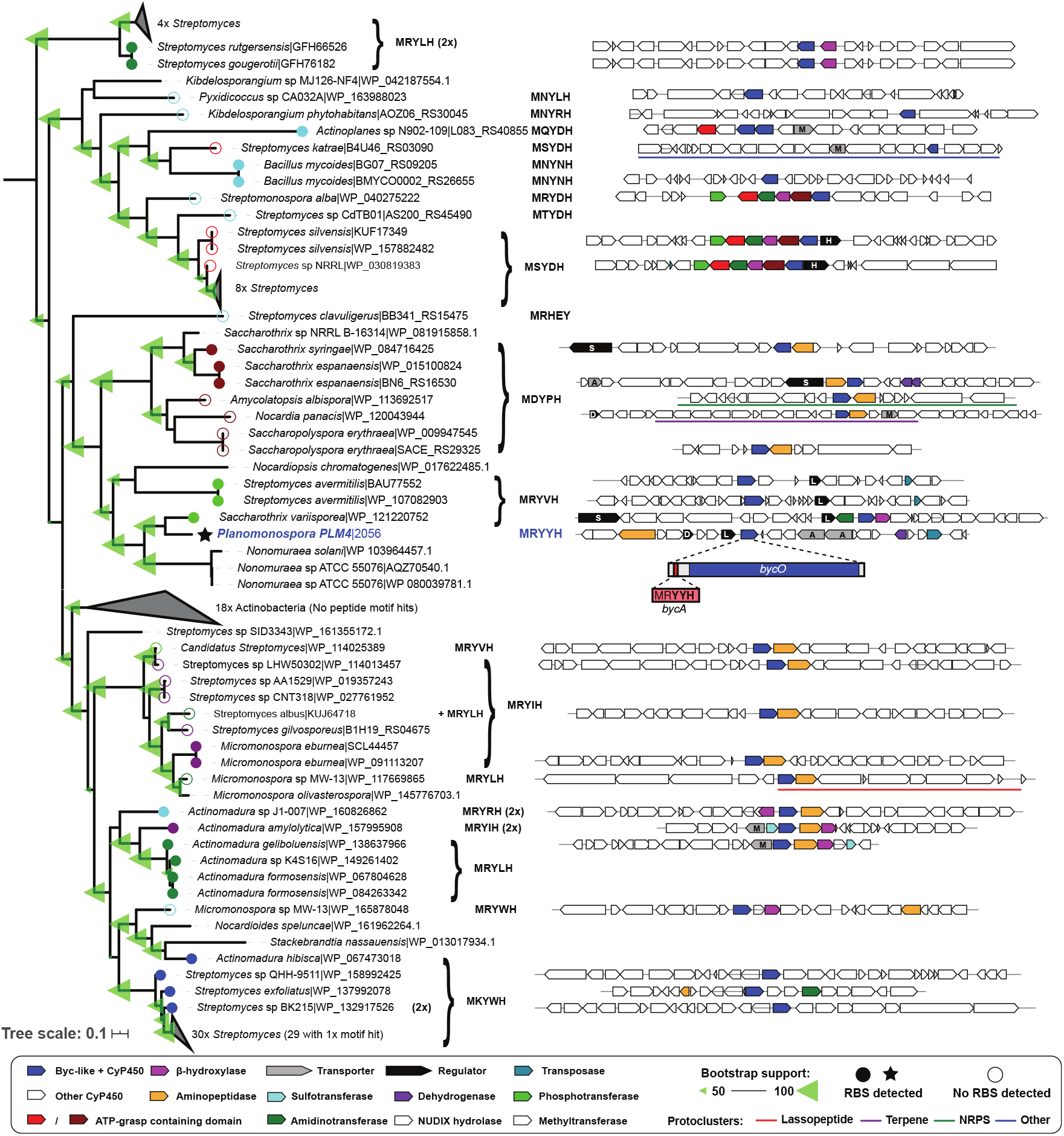
Summary phylogeny of cytochrome P450s, likely associated with biarylicin biosynthesis. Identical pentapeptide motif hits are shown as colored circles corresponding to the motif printed to the right of the tree. Filled and unfilled circles indicate those with and without a detected Shine-Dalgarno RBS, respectively. Selected regions of *byc*-like clusters are shown to the right of motifs, illustrating a variety of contexts and shared coding sequences potentially linked to the minimal P450 cluster (highlighted in blue). Expanded clades and detailed annotations are accessible at: https://itol.embl.de/tree/2131271061176991591098016.

We propose the name biarylicins for this family of *N*-acetylated tripeptides, carrying a crosslink between the aromatic side chains of amino acids 1 and 3. Accordingly, compounds **1A**, **1B**, K-13, OF-4949 and pseudosporamide would become biarylicins YYH, YFH, YYY, YNY and YPW, respectively. Since the first comprehensive literature overview of just a few years ago,^2^ several new RiPP families have been uncovered, mostly through searching precursor peptide-related features in genome sequences.^17^ RiPPs such as the biarylicins can however not be detected by current bioinformatic search strategies because of the extremely small size of the *bycA* gene. Nonetheless, as shown herein, with hindsight from genomic data, these minimal RIPPs can be identified via cytochome P450 homology and screening for the short peptide sequences in many bacterial genomes. While occurring mostly in actinobacteria, one such BGC was also detected in a *Pyxidicoccus* strain. It remains to be determined whether the search criteria implemented in this work (the pentapeptide MxYxY/H-encoding gene within ± 500 bp from a cytochrome P450 monooxygenase gene) were too narrow to detect all similar BGCs.

While the *byc* BGC is extremely small, it poses nonetheless some biochemical challenges for the specificity of the biaryl cross-link. The basic amino acid, often found at position −1, might act as a signal for processing. Also, ribosomal binding sites other than Shine-Dalgarno sequences might precede the peptides. Further studies will be needed to understand the timing and sequence requirements for installing a cross-link and leader peptide cleavage. Since the biaryl cross-link imparts proteolytic stability, understanding its specificity might have important applications in stabilizing peptides for different applications.

In conclusion, after identifying cross-linked, cyclic tripeptides produced by different strains of *Planomonospora*, we uncovered a new, widespread family of RiPPs specified by a minimal, two-gene BGC encoding a pentapeptide, so far the smallest gene reported, and a cytochrome P450 monooxygenase. Our understanding of the biological activities of biarylicins is extremely limited. The different protease inhibitory activities seen with **1A**, **2** and **3** suggest that the amino acid sequence and/or ring size can strongly influence bioactivity. Thus, their biological properties and their role in the producing organisms await discovery.

## Experimental Procedures

### Isolation and purification of compound 1A

Chemicals and molecular biological agents were purchased from standard commercial sources. General conditions for strain cultivations were as described elsewhere.^4^ For isolation and purification of compound **1A**, 1.5 mL of a frozen stock of strain ID82291 was used to inoculate 15mL AF medium in a 50mL baffled flask.^18^ After 72 hours, 10mL were used to inoculate 100mL fresh AF medium in a 500mL baffled flask. After further 72 hours, 100mL was used to inoculate 900mL AF medium in a 3000mL flask, containing 50g 5 mm glass beads. After 7 days, the culture supernatant was cleared by both centrifugation and filtration and the cleared broth was adsorbed on 200mL HP20 resin (Diaion). The loaded resin was washed with demineralized water and eluted with aqueous 50% MeOH (V/V). The eluate was brought to dryness. The dried extract was taken up in 10% acetonitrile/90% water and pre-fractionated on a Biotage SNAP Ultra HP-Sphere C18 25μm 12g with a CombiFlash system (Teledyne ISCO, Nebraska, USA). Phases A and B were water (MilliQ, Merck) and acetonitrile (Sigma-Aldrich), respectively. A gradient with steps of 5, 5, 30, 100 and 100 % phase B at 0, 5, 20, 25 and 30 min, respectively, was applied. After screening fractions by LC-MS, **1A**-containing fractions were dried, taken up in 0.05% trifluoroacetic acid (TFA) in 10% MeCN/90% MilliQ water and processed further by semi-preparative HPLC on a Shimadzu LC-2010CHT instrument, equipped with a 250/10 Nucleodur C18 Pyramid 5M column (Macherey-Nagel) and a C18 pre-column. An isocratic method with 10% phase B (A: 0.05% TFA in MilliQ water, pH 2.5; B: MeCN) at 3mL/min, column temperature 40°C was used for purification. Due to co-eluting contaminants, the semipurified substance was purified for a second time under the same conditions, except for the change of phase A to MilliQ water. Drying of the eluent yielded 5 mg of a white solid.

### Analytical procedures

High-resolution MS spectra acquisition was performed on a Dionex UltiMate 3000 (Thermo Scientific) coupled with a micrOTOF-Q III (Bruker) instrument equipped with an electrospray interface (ESI), using an Atlantis T3 C_18_ 5μm x 4.6 mm x 50 mm column, as reported elsewhere.^4^ MS measurements for monitoring metabolite production were performed on a Dionex UltiMate 3000 (Thermo Scientific) coupled with an LCQ Fleet (Thermo Scientific) mass spectrometer equipped with an electrospray interface (ESI) on an Atlantis T3 C_18_ 5μm x 4.6 mm x 50 mm column, as reported elsewhere.^19^ MS measurements for analysing **1A** after acid hydrolysis were performed on the same instrument, applying a hydrophilic gradient with 0, 0, 25, 95, 95, 0 and 0% phase B (MeCN) at 0, 1, 7, 8, 12, 12.5 and 14 min, respectively, with the *m/z* range set to 110–1000 *m/z.* Phase A was 0.05% TFA in MilliQ water. UV-VIS signals (190-600 nm) were acquired using the built-in diode array detector of the Dionex UltiMate 3000 (Thermo Scientific). Mono-and bidimensional NMR spectra were measured in DMSO-*d_6_* or CD_3_OD at 298K using a Bruker Avance 300 MHz spectrometer.

### Hydrolysis of 1A

A small amount of purified compound **1A** (< 1mg) was dissolved in 250μL 0.1M sodium borate buffer (pH 9), followed by addition of 250μL fresh 12M HCl (Sigma-Aldrich). The solution was heated to 120°C and sampled at 24, 48 and 96 hours. The samples were analysed by LC-MS analysis, using a hydrophilic gradient as described above.

### 1A production curve

1.5 mL of a frozen stock culture of strain ID82291 was used to inoculate 15 mL AF medium in a 50 mL baffled flask. After 72 hours, 1.5 mL were used to inoculate each 15 mL AF medium, 15 mL RARE3 medium^20^ or 15 mL AF medium with 30% D_2_O (v/v) in 50 mL baffled flasks and further cultivated under identical conditions. Production of **1A** was monitored by sampling 1 mL culture every 24 hours. Samples were centrifuged and the supernatant, after adsorption on 200 μL of HP20 resin (Diaion) for 1 hour, was eluted with 200 μL of methanol (Sigma). Samples were analysed by LC-MS as described above.

### Biarylicin YYH (1A)

white crystalline powder; [α]⊓ = −165 (c 0.11, H_2_O pH10, 25°C); 1H and 13C NMR, Table in Supporting Information; HRESIMS *m/z* 522.198 [M+H]^+^ (calcd for C_26_H_27_N_5_O_7_, 522.1988); ESI-MS/MS ([M+H]^+^) *m/z* (%) 269.139 (100), 359.134 (84), 315.143 (69), 494.201 (66), 289.128 (61), 285.133 (56), 331.140 (54), 243.122 (51), 313.129 (37), 448.198 (27), 476.191 (17), 228.111 (10), 406.181 (9), 214.094 (9), 199.081 (7).

### Cloning and heterologous Expression of *bycAO*

A DNA stretch containing *bycAO* (1303 bp) was cloned into pSET152_*ermE\** via Gibson assembly. Linear DNA fragments were generated by PCR using primers designed to include 20 bp overlap sequences (pSET_fwd: CCTCTC-TAGAGTCGACCTGCAGC, pSET_rev: CCTTCCGTACCTCCGTTGCT, *bycAO*_fwd: AGCAACG-GAGGTACGGAAGGAGGAGGTGTGCGATGCGC and *bycAO*_rev: GCAGGTCGACTCTAGA-GAGGCTAGCGGGGGAGAAGGAC). Purified linear vector and insert were mixed (ratio 1:3) with 15 μL of Gibson assembly master mixture (1X ISO buffer, 10 U/μL T5 exonuclease, 2 U/μL Phusion polymerase, and 40 U/μL Taq ligase) and incubated at 50 °C for 1 hour. 5 μL of reaction were transformed into chemically competent *E. coli* DH5α cells and plated onto LB agar supplemented with 50 μg/mL apramycin. Positive clones containing the plasmid pSET-*bycAO* were selected by colony PCR (primers: pSET-*bycAO*_fwd: GCGTCGATTTTTGTGATGCTCG and pSET-bycAO_rev: CAGCGAATTCGGAAAACGGC) and confirmed using restriction digest and sequencing. pSET*bycAO* was conjugated into *S. coelicolor* M1152 using a standard intergeneric conjugation protocol with the methylation-deficient *E. coli* ET12567/pUZ8002/pSET-*bycAO* as the donor. Cells were spread onto MS agar (20 g/L soya flour, 20 g/L mannitol, 20 g/L agar, 50 mM MgCl_2_) and incubated at 30 °C. After 20 hrs, 0.5 mg nalidixic acid and 1 mg apramycin were overlaid on the plates and incubation continued until ex-conjugants showed up. Single colonies were spread on new plates containing 25 mg/L of nalidixic acid and 50 mg/L of apramycin to confirm the ex-conjugants. Colonies containing *bycAO* were validated by colony PCR and cultivated in TSB broth for 12 days at 30 °C. Cultures were extracted as described above and analyzed by HR-LC-MS/MS.

### Phylogenetic analyses

A set of complete genomes from the antiSMASH database were used along with top blast results to PLM4_2056 from *Planomonospora* sp. ID82291 (NCBI reference JABTEX000000000). Cytochrome P450 enzymes were identified via a hidden Markov model search using the Pfam (PF00067) P450 model and trusted bit-score cutoffs. After ensuring model and query alignment coverage over 70%, similar sequences (>95% amino acid ID) were clustered via an all vs. all blast and one representative was used for the final dataset, which yielded over 3,300 sequences. Any hit to the motif search analysis (see below) were also included in this dataset. All sequences were aligned via MAFFT using the local iterative (-linsi) option followed by alignment trimming using the trimal application with automated option to improve maximum-likelihood tree inference. Due to the large number of sequences FastTree was used to infer the final phylogenetic gene tree.

### Peptide motif search

A custom python script (https://git.wur.nl/snippets/65) was used to examine 6-frame translations of corresponding genomic regions +/-500bp of identified cytochrome P450 enzymes. This reported any motif matching MxYx[Y/H]*, where x is any amino acid and * is a stop codon. Reported positions were then cross-referenced to highlight motifs occurring up/downstream of the P450 CDS. An additional screen of these results for the Shine-Dalgarno sequence AGGAGG was preformed upstream of all motif positions to annotate RBS sites. Identified peptides were mapped onto the generated, P450-based phylogenetic tree and visualized using the iTOL-webserver. ^21,22^

## Acknowledgement

The authors thank Stefan Kehraus for measuring optical rotation of **1A**. This work has received funding from the European Union’s Horizon 2020 research and innovation program under grant agreement No.721484 (Train2Target). M.G.S.Z. acknowledges the DAAD for a Ph.D scholarship. M.C. received funding from the German Research Foundation (DFG) grant nr. FOR2372. M.A. and M.H.M. were supported by an ERA NET CoBiotech (Bestbiosurf) grant through the Netherlands Organization for Scientific Research (NWO) [053.80.739].

## Supporting Information Available

The following files are available free of charge.

- Supporting Information: Figures S1-S21.

