## Supporting Information for "A biaryl-linked tripeptide from *Planomonospora* leads to widespread class of minimal RiPP gene clusters"

21 NMR experiment: combined NOESY-TOCSY of biarylicin A in DMSO-*d6* . 23

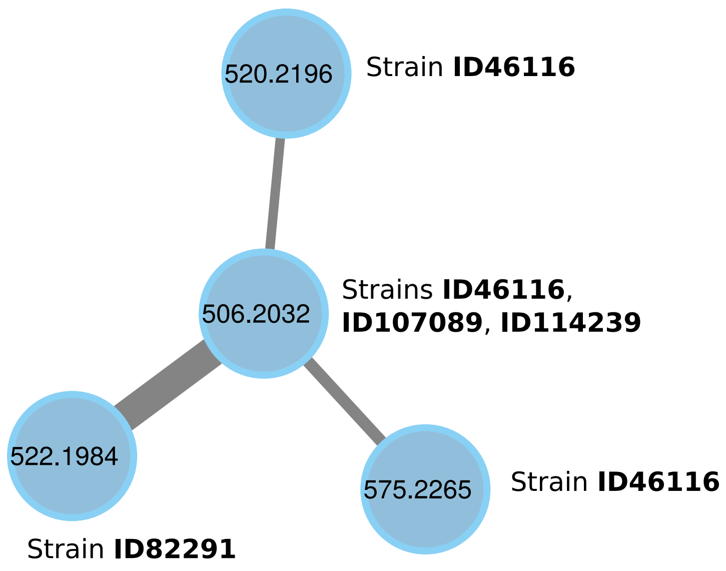

Figure 1: Molecular family showing the occurrence of biarylicin A (522.19) and B (506.2), as well as two minor, uncharacterized congeners detected only in strain ID46116.

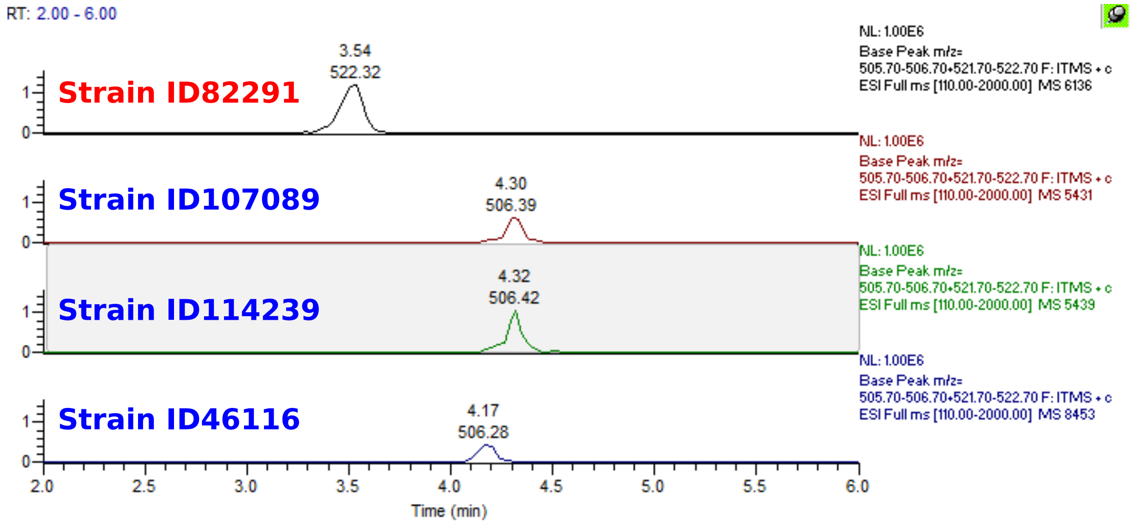

Figure 2: LCMS-traces showing the mutually exclusive occurrence of biarylicin A ($m/z$ 522, red) and biarylicin B ($m/z$ 506, blue). For strain ID46116, the slight difference in comparison to ID107089 and ID114239 can be explained by retention time shift.

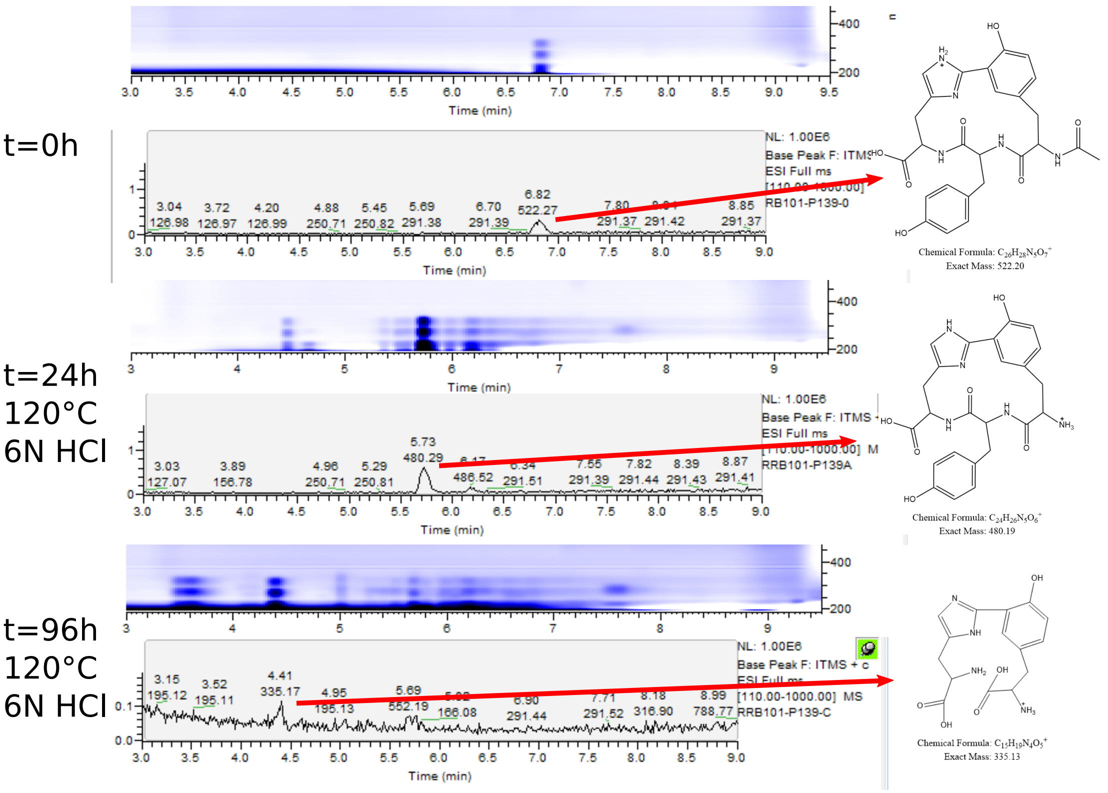

Figure 3: LCMS-traces of hydrolysis of biarylicin A ($m/z$ 522), using the hydrophilic LCMS-method. First, deacetylation ($m/z$ 480) can be observed, while eventually, hydrolysis and loss of the central tyrosine ($m/z$ 335) occurs.

**A) $MS^2$-fragmentation pattern**

| **522.1982 *m/z*[a]** $C_{26}H_{28}N_5O_7$ | **506.2029 *m/z*[b]** $C_{26}H_{28}N_5O_6$ |
|---|---|
| 504.1862 (-18.01) | 488.1983 (-18.00) |
| 494.2009 (-27.99) | 478.2077 (-27.99) |
| 476.1909 (-46.01) | 460.1964 (-46.01) |
| 480.1853 (-42.01) | 464.1918 (-42.01) |
| 448.1979 (-74.00) | 432.2001 (-74.00) |
| 359.1339 (-163.06) | 359.1339 (-147.07) |
| 331.1400 | 331.1391 |
| 315.1432 | 315.1441 |
| 313.1290 | 313.1267 |
| 289.1277 | 289.1282 |
| 285.1327 | 285.1319 |
| 269.1388 | 269.1398 |
| 243.1217 | 243.1203 |
| 228.1113 | 228.1100 |
| 214.0943 | 214.0943 |
| 199.0812 | 199.0846 |

[a]Biarylicin A
[b]Biarylicin B

**B)**

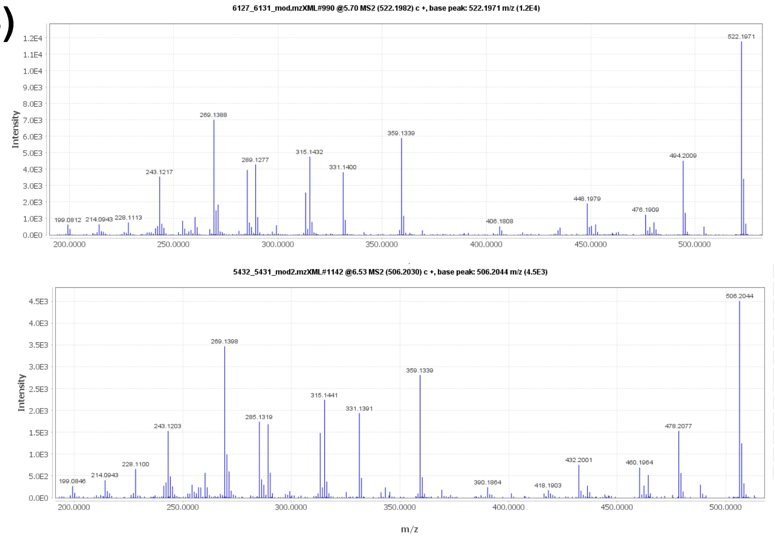

Figure 4: $MS^2$-fragmentation patterns of biarylicins, investigated in this study. (A) compares the main fragments of both biarylicin A and B, with the only difference between them indicating a phenylalanine instead of a tyrosine loss. (B) shows original HR-MS/MS spectra for biarylicin A (top) and B (bottom).

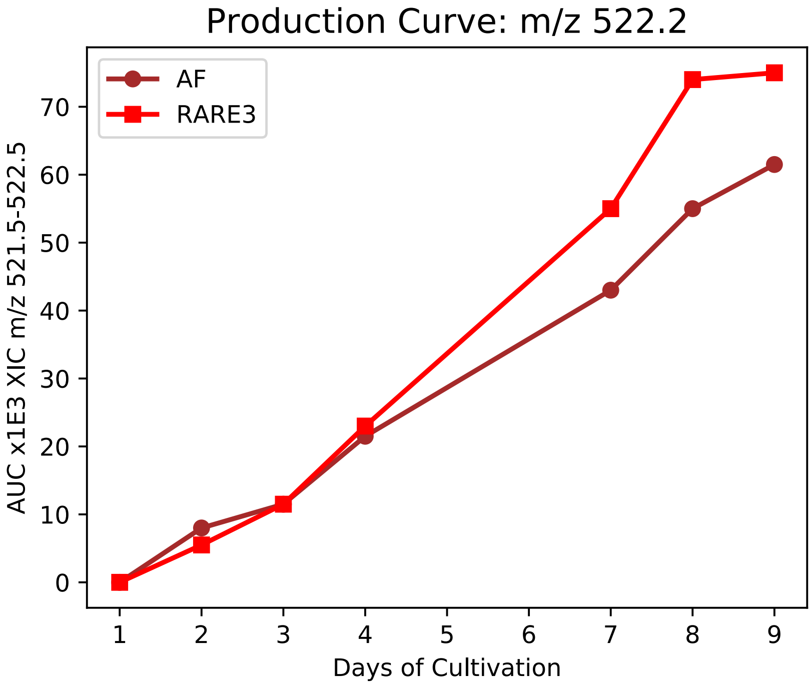

Figure 5: Visualization of biarylicins A production by *Planomonospora* strain ID82291 in two different media over time, detected by LCMS-analysis. The y-axis gives the area under the curve of the extracted ion chromatogram for $m/z$ 522.

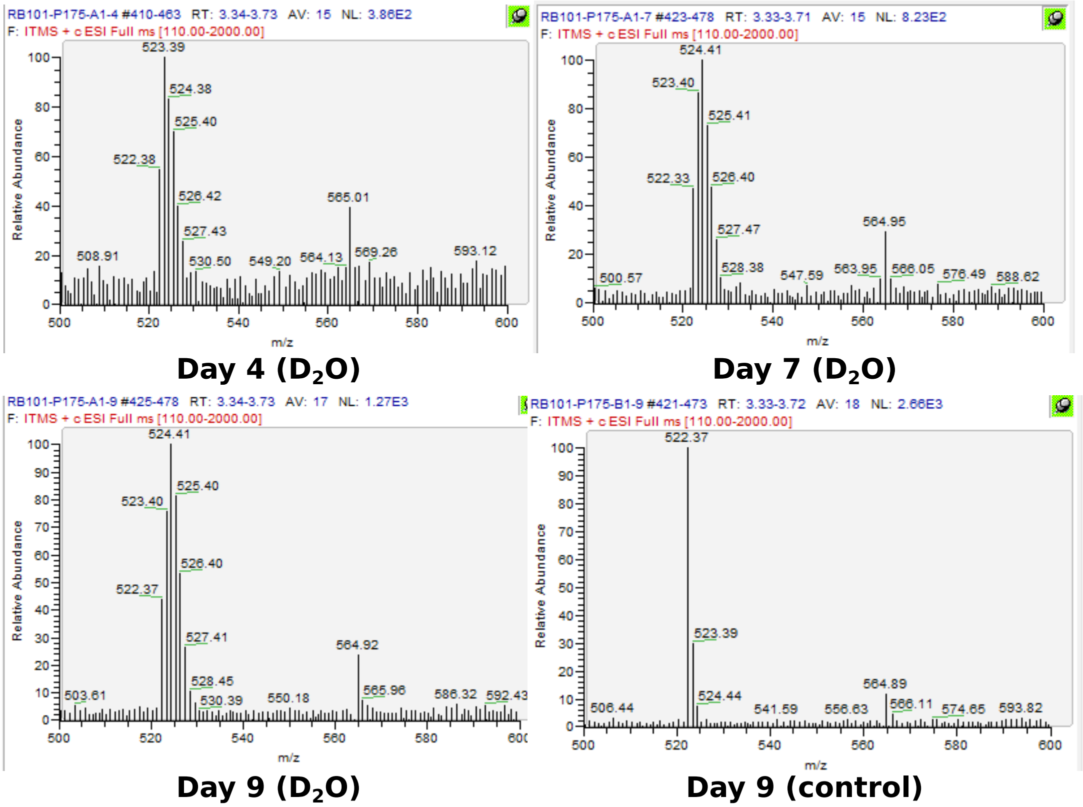

Figure 6: Cultivation of strain ID82291 in $D_2O$-supplemented medium leads to incorporation of deuterium into biarylicins A, resulting in Gaussian isotope patterns in LCMS. In the deuterium-free control, a normal, monotonic isotope pattern can be seen.

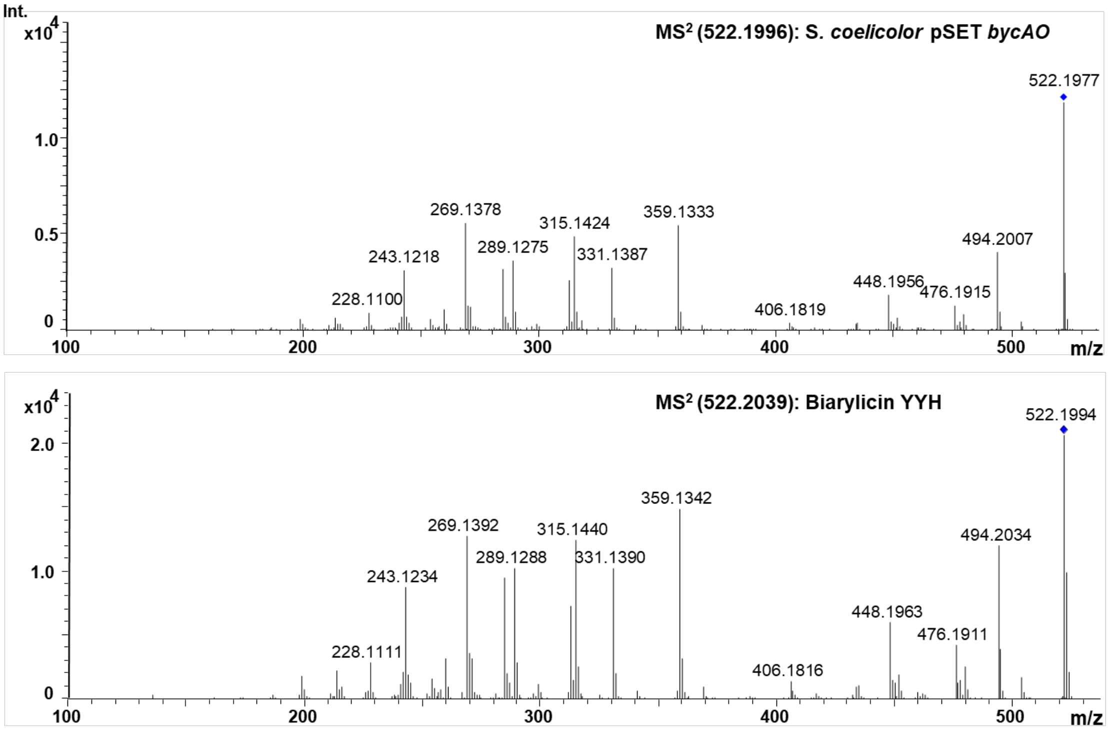

Figure 7: Comparision of tandem mass fragmentation spectra of (A) biarylicin YYH from heterologous expression in *Streptomyces coelicolor* M1152 and (B) authentic standard of biarylicin YYH, isolated from *Planomonospora* sp. ID82291.

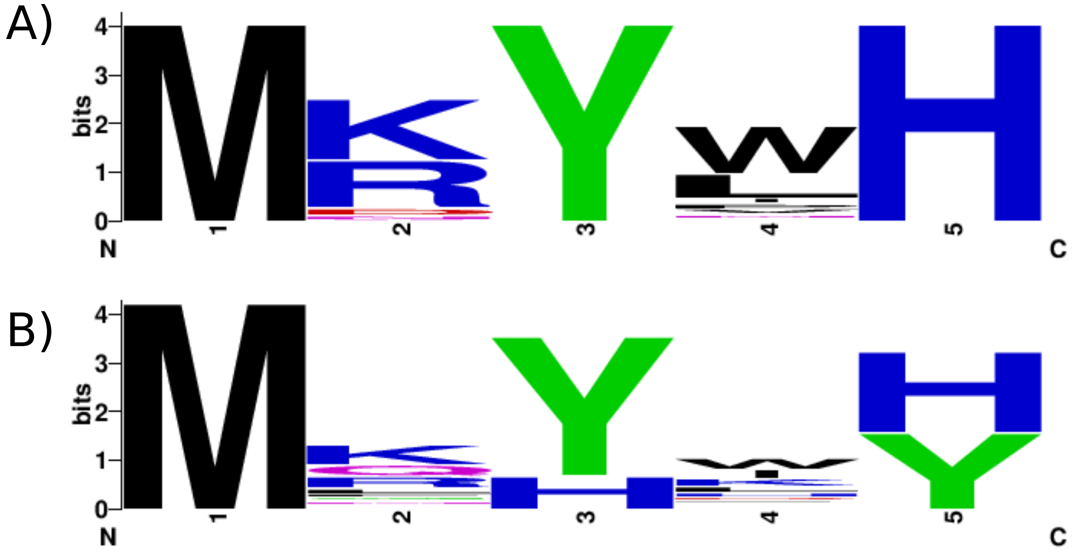

Figure 8: Amino acid distribution in detected pentapeptides. (A) Distribution of amino acids in pentapeptides, preceded by a Shine-Dalgarno ribosomal binding site. (B) Distribution in all pentapeptides.

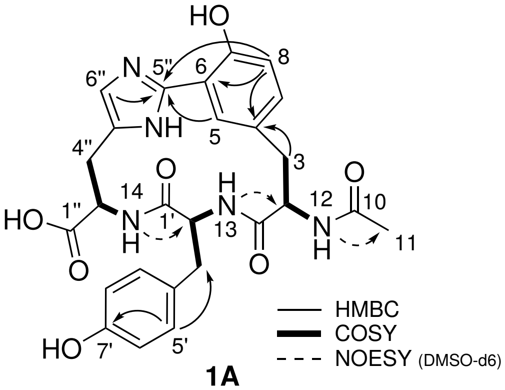

**1A**

| **Position** | **biarylicin YYH (1A)** $\delta_C$ **(type) DMSO-*d6*** | $\delta_H$ **DMSO-*d6*** | $\delta_C$ **(type)** $CD_3OD$ | $\delta_H$ $CD_3OD$ | **COSY**[a] | **NOESY**[a] | **HMBC**[a] |
|---|---|---|---|---|---|---|---|
| 1 | 172 (C) | - | 173.5 (C) | - | - | - | - |
| 2 | 54 (CH) | 4.6 | 55.5 (CH) | 4.31 | **3**, 12 | - | *1, 3, 4* |
| 3 | 37 ($CH_2$) | 2.51-3.12 | 37.2 ($CH_2$) | 2.66-3.24 | **2** | - | **1**, *2*, **4** |
| 4 | 130 (C) | - | 130.3 (C) | - | - | - | - |
| 5 | - | - | 128.7 (CH) | 7.32 | - | - | *3, 7, 5''* |
| 6 | - | - | 110 (C) | - | - | - | - |
| 7 | - | - | 155 (C) | - | - | - | - |
| 8 | 117 (C) | 6.8 | 117 (CH) | 6.97 | *9* | - | *3*, **4**, *6, 5''* |
| 9 | - | - | 136 (CH) | 7.27 | *8* | - | *3, 7* |
| 10 | 170.5 (C) | - | 171.9 (C) | - | - | - | - |
| 11 | 24 ($CH_3$) | 1.78 | 22.3 ($CH_3$) | 1.9 | - | - | **10** |
| 1' | - | - | 173.1 (C) | - | - | - | - |
| 2' | 56 (CH) | 4.7 | 55.7 (CH) | 4.92 | **3'**, 13 | - | *3', 4', 1'* |
| 3' | 39 (CH2) | 2.7-2.98 | 39.4 ($CH_2$) | 2.87-2.66 | **2'** | - | *1', 2', 4', 5', 9'* |
| 4' | - | - | 128 (C) | - | - | - | - |
| 5'/9' | 132 (CH) | 6.97 | 131 (CH) | 6.98 | 6', 8' | - | **3'** ,*7'*, *9'* |
| 6'/8' | 116 (CH) | 6.58 | 116 (CH) | 6.63 | 5', 9' | - | *9'* |
| 7' | 156 (C) | - | 157 (C) | - | - | - | - |
| 1'' | - | - | 172.3 (C) | - | - | - | - |
| 2'' | 55 (CH) | 4.49 | 54 (CH) | 4.48 | **3''**, 14 | - | *1'', 3'', 4''* |
| 3'' | 29.5 ($CH_2$) | 2.94 | 29 ($CH_2$) | 2.77-3.53 | **2''** | - | *2'', 4'', 6''* |
| 4'' | - | - | 131.4 (C) | - | - | - | - |
| 5'' | - | - | 142 (C) | - | - | - | - |
| 6'' | - | - | 117 (CH) | 7.35 | - | - | *2'', 3'', 5''* |
| NH(12) | - | 7.83 | - | - | 2 | 11 | 10 |
| NH(13) | - | 8.38 | - | - | 2' | 2 | - |
| NH(14) | - | 9.5 | - | - | 2'' | 2' | - |

[a]2D correlations: **boldface**=both $CD_3OD$ and DMSO-*d6*; *italics*=$CD_3OD$; normal=DMSO-*d6*

Figure 9: NMR spectroscopic data for **1A**, recorded in $CD_3OD$ and DMSO-*d6* (300 MHz).

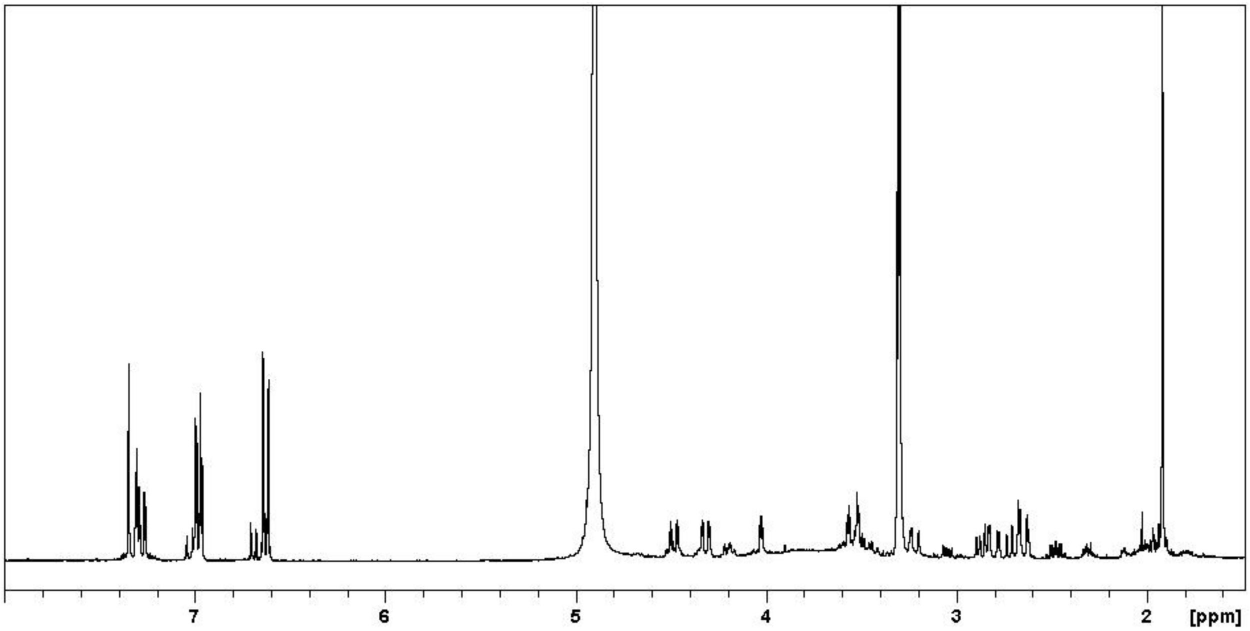

Figure 10: $^{1}H$ of biarylicin A in $CD_3OD$.

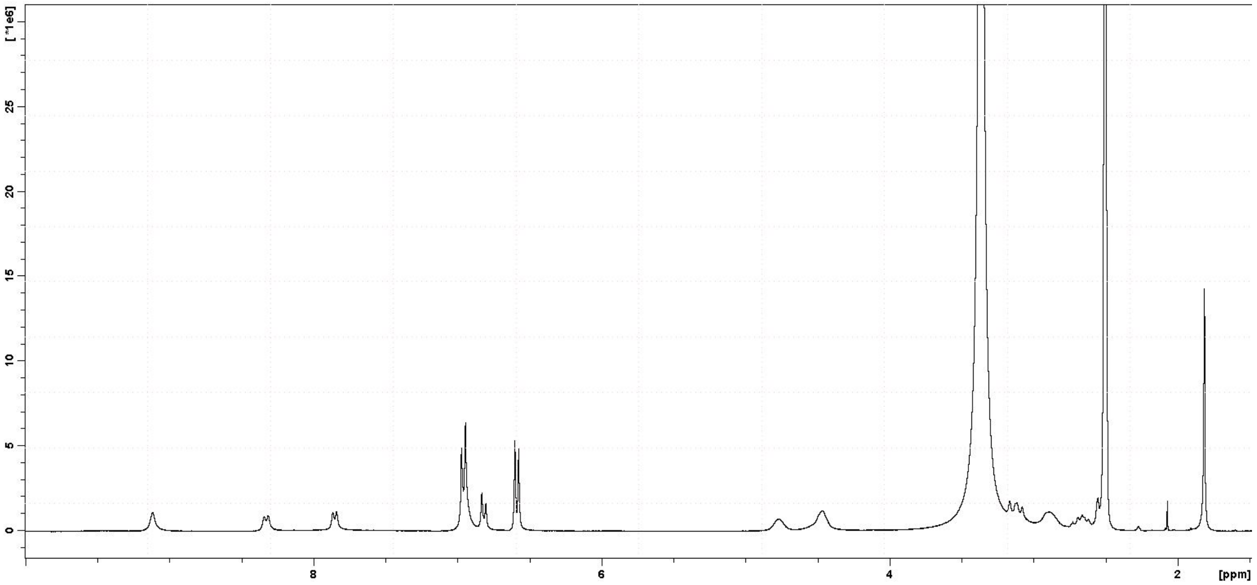

Figure 11: $^{1}H$ of biarylicin A in DMSO-*d6*.

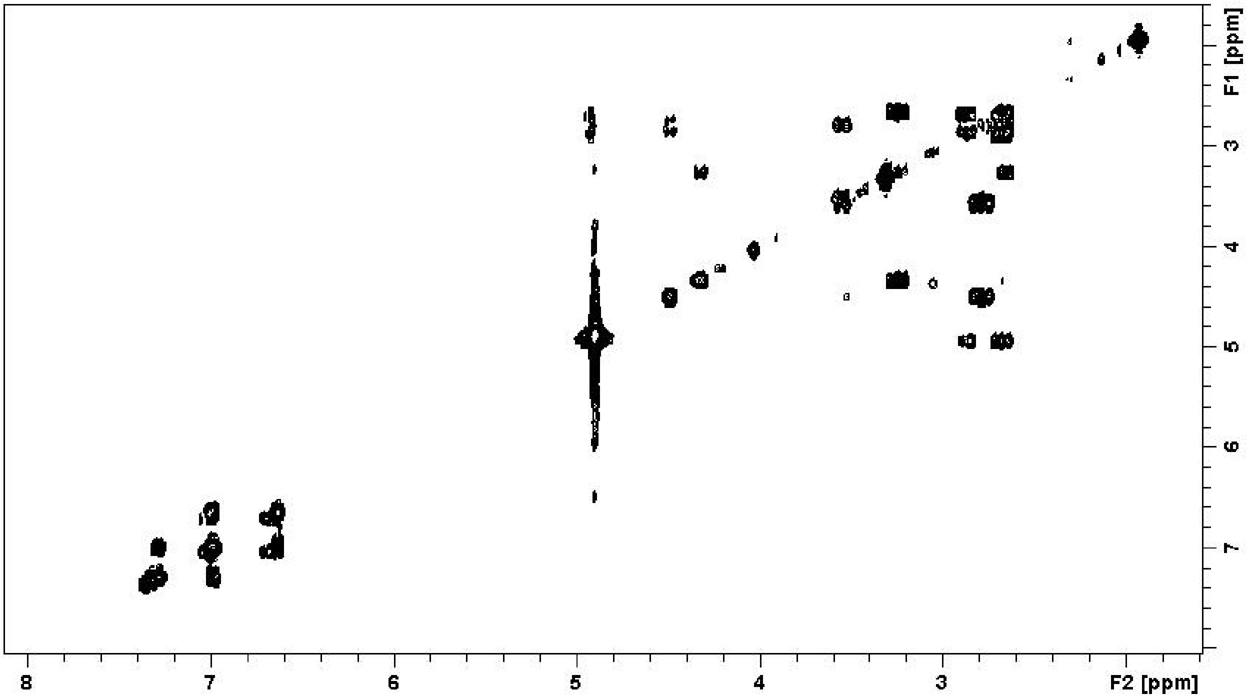

Figure 12: COSY of biarylicin A in $CD_3OD$.

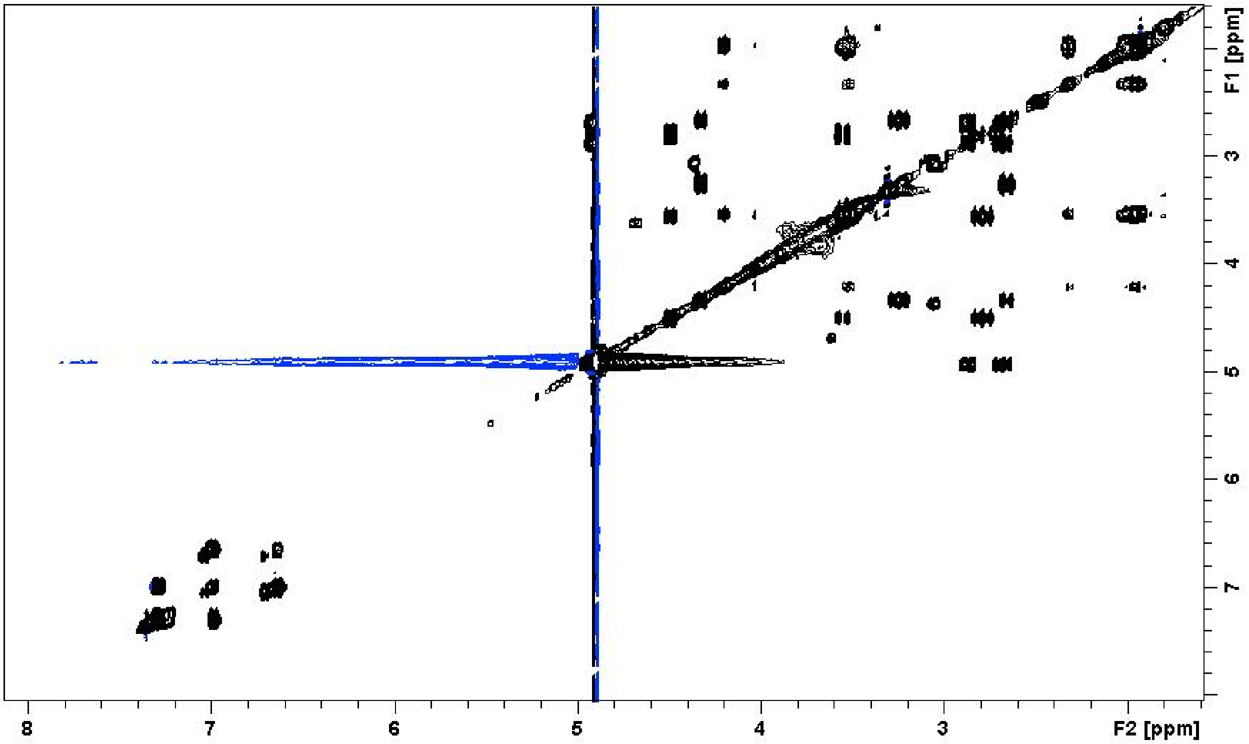

Figure 13: TOCSY of biarylicin A in $CD_3OD$.

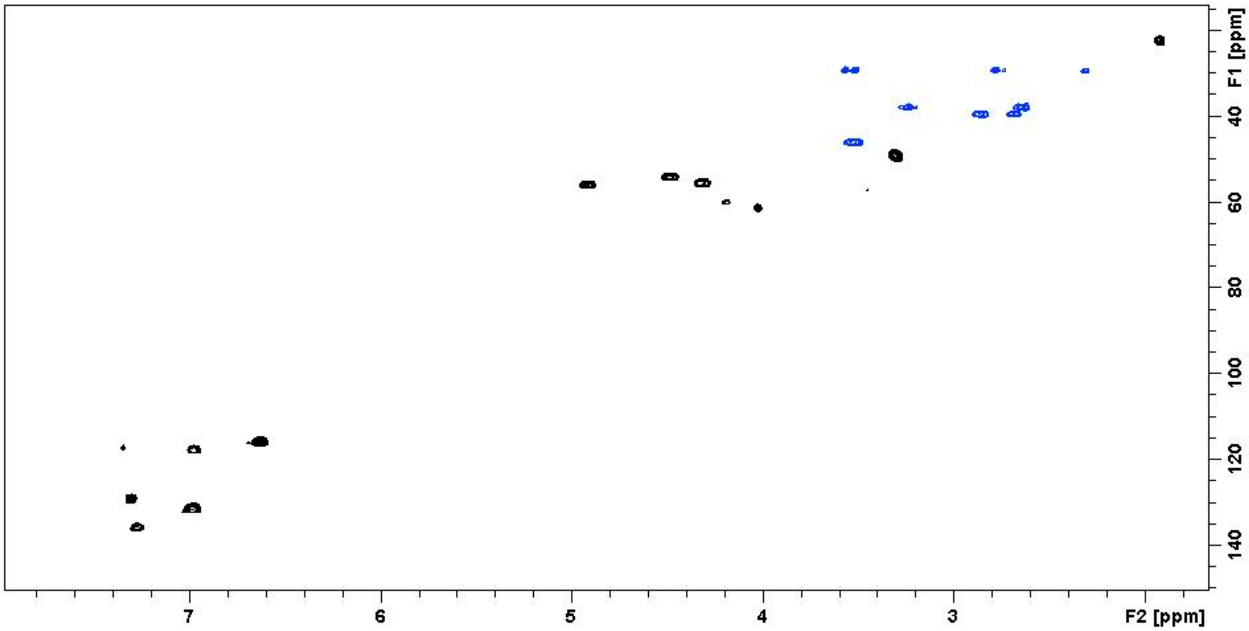

Figure 14: HSQC of biarylicin A in $CD_3OD$.

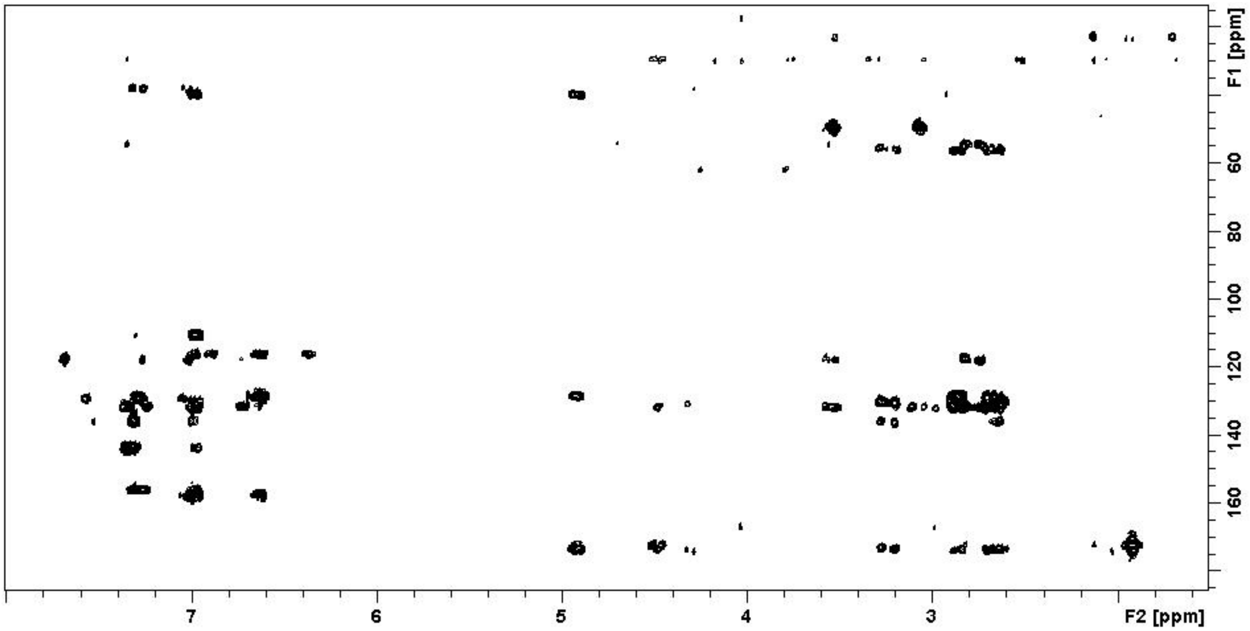

Figure 15: HMBC of biarylicin A in $CD_3OD$.

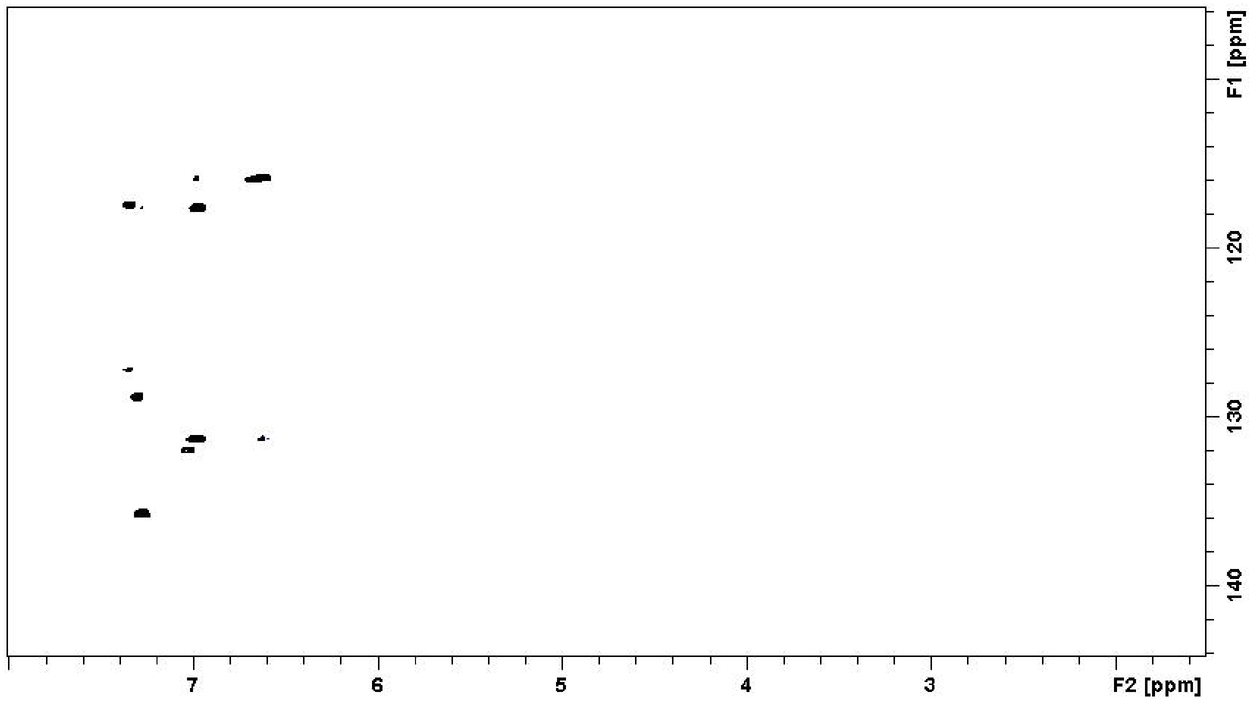

Figure 16: Selective band HSQC of biarylicin A in $CD_3OD$.

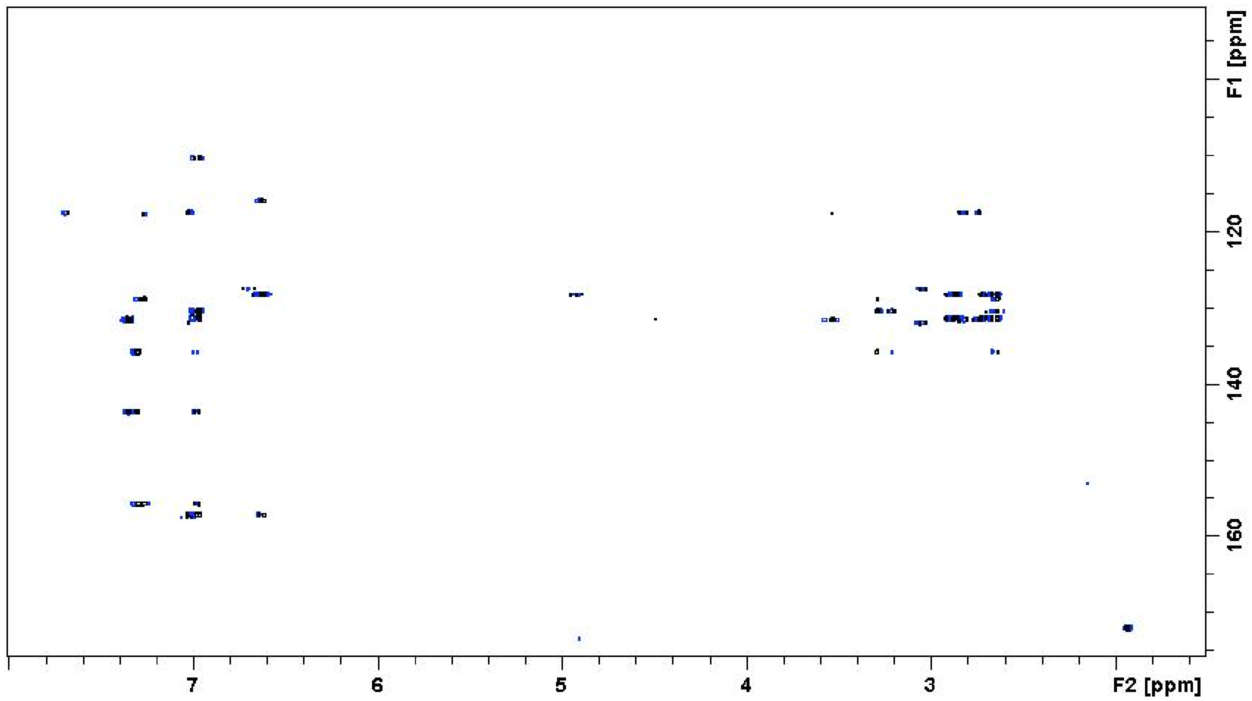

Figure 17: Selective band HMBC of biarylicin A in $CD_3OD$.

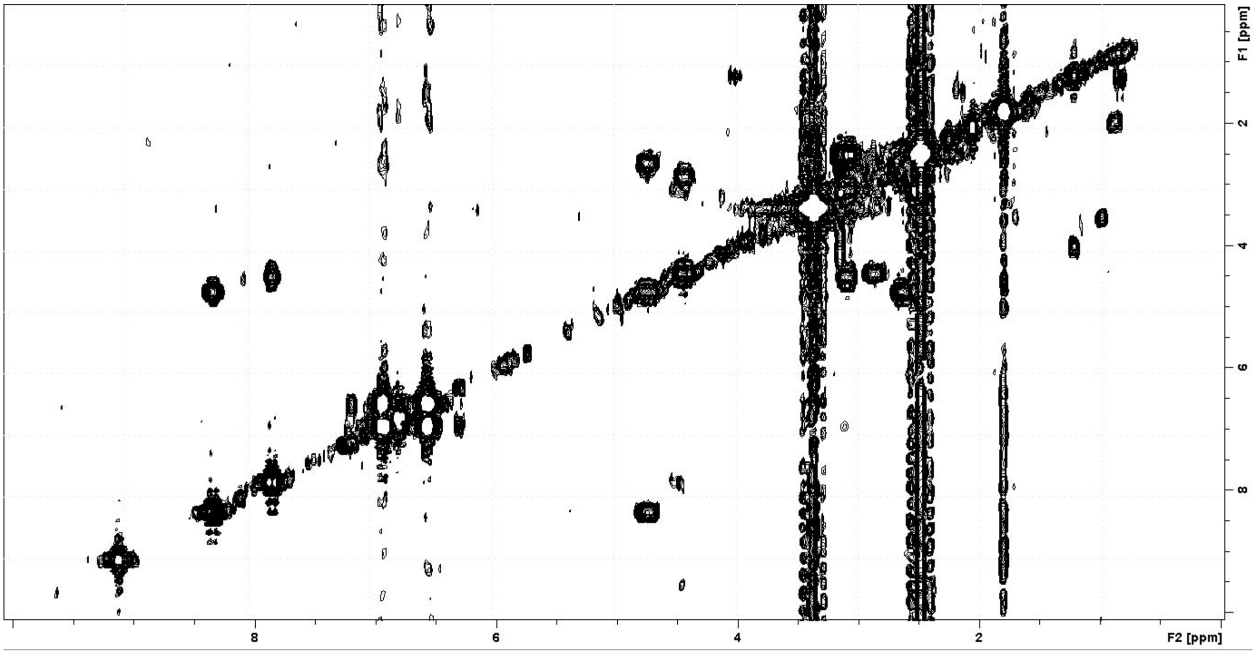

Figure 18: COSY of biarylicin A in DMSO-*d6*.

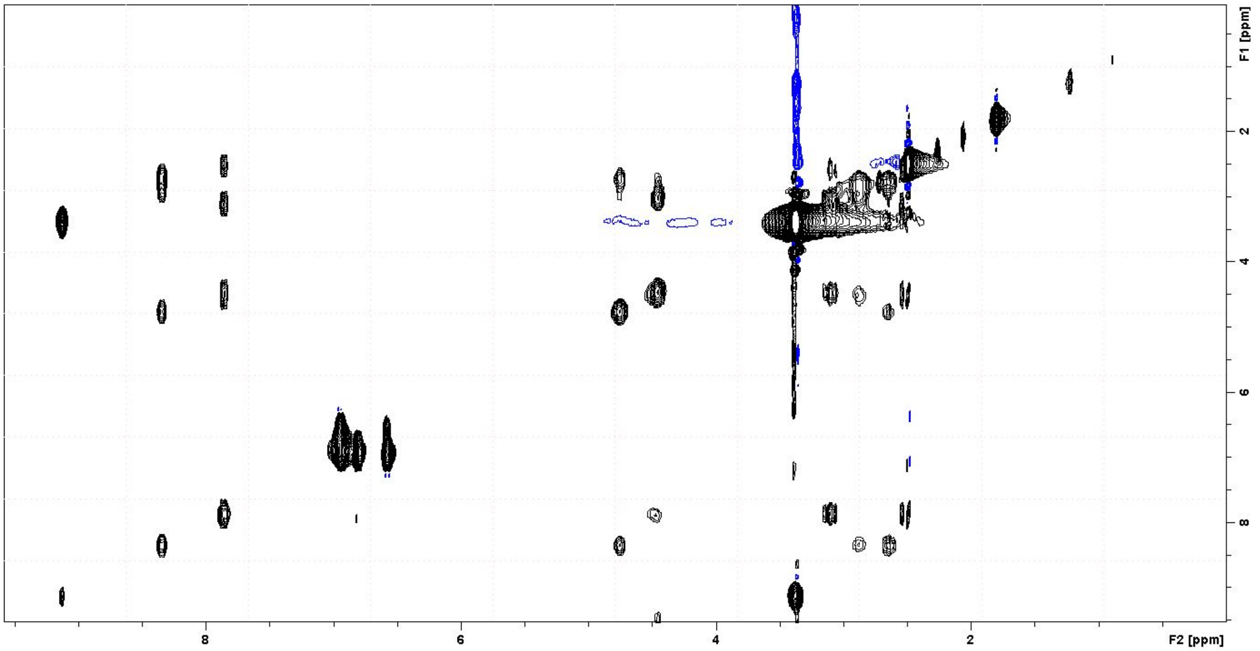

Figure 19: TOCSY of biarylicin A in DMSO-*d6*.

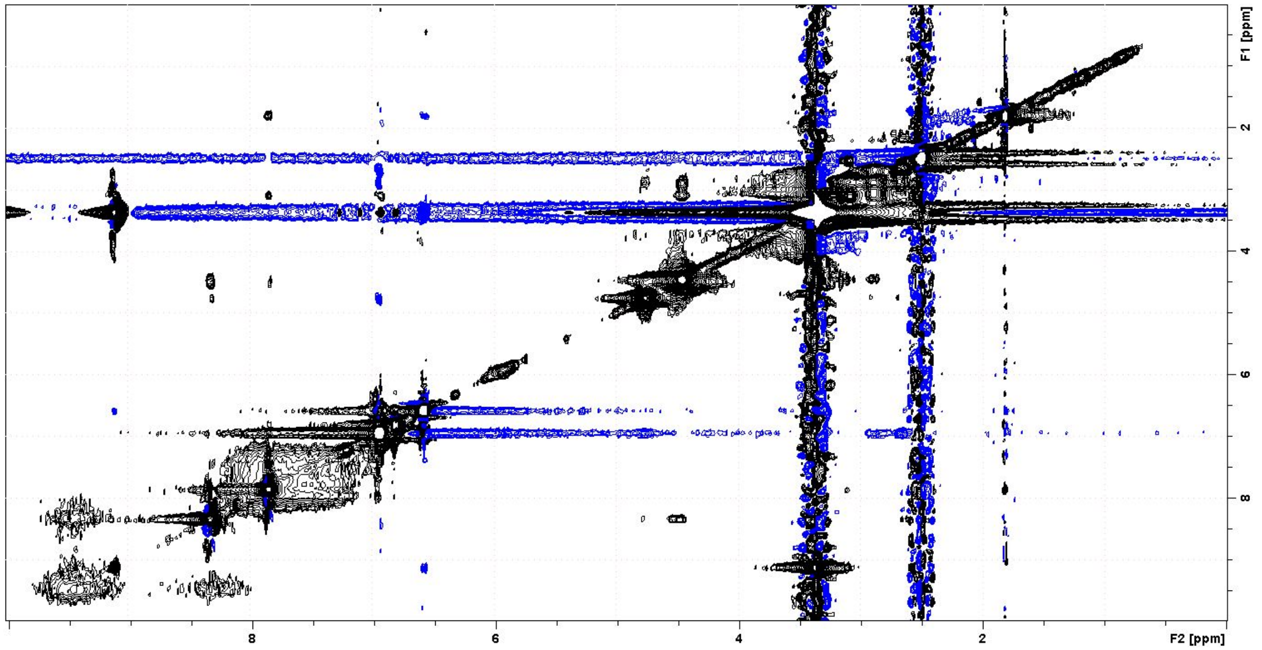

Figure 20: NOESY of biarylicin A in DMSO-*d6*.

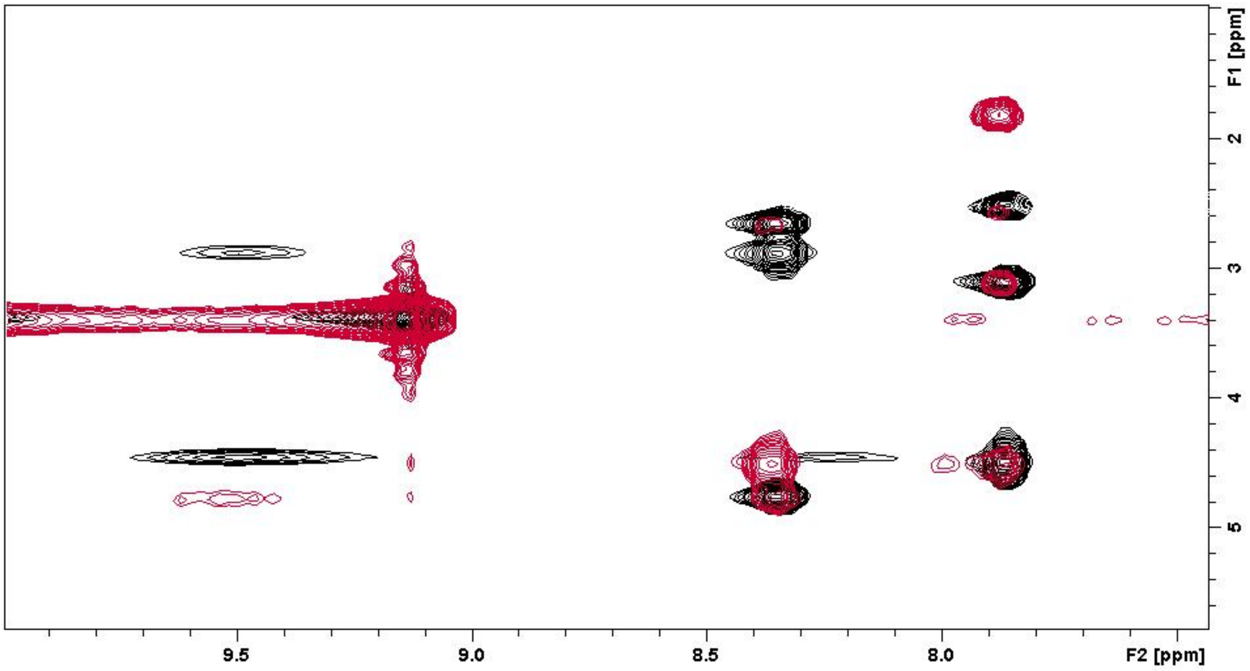

Figure 21: Zoom into the amidic region in a combined NOESY-TOCSY experiment of biarylicin A in DMSO-*d6*.
